# A single point mutation in a TssB/VipA homolog disrupts sheath formation in the type VI secretion system of *Proteus mirabilis*

**DOI:** 10.1101/152124

**Authors:** Christina C. Saak, Martha A. Zepeda-Rivera, Karine A. Gibbs

## Abstract

The type VI secretion (T6S) system is a molecular device for the delivery of proteins from one cell into another. T6S function depends on the contractile sheath comprised of TssB/VipA and TssC/VipB proteins. We previously reported on a mutant variant of TssB that disrupts T6S-dependent export of the self-identity protein, IdsD, in the bacterium *Proteus mirabilis*. Here we determined the mechanism underlying that initial observation. We show that T6S-dependent export of multiple self-recognition proteins is abrogated in this mutant strain. We have mapped the mutation, which is a single amino acid change, to a region predicted to be involved in the formation of the TssB-TssC sheath. We have demonstrated that this mutation does indeed inhibit sheath formation, thereby explaining the global disruption of T6S activity. We propose that this mutation could be utilized as an important tool for studying functions and behaviors associated with T6S systems.

## Introduction

Type VI secretion (T6S) systems, widely found among Gram-negative bacteria, are cell-puncturing devices that deliver cargo proteins from one cell into another or into the environment (1-22). T6S function, and ultimately cargo delivery, depends on a contractile sheath made up of TssB/VipA (Pfam family PF05591) and TssC/VipB (Pfam family PF05943) proteins (6, 23, 24). TssB and TssC bind one another to form protomers, which in turn assemble into the contractile sheath (25-27). Subcellular visualization of the T6S machinery has relied on the fusion of TssB to fluorescent proteins such as superfolder Green Fluorescent Protein (sfGFP) and mCherry, thereby allowing the detection of sheath dynamics and interactions between TssB and TssC, as well as measuring the active firing of T6S machines within a population (4-6, 17, 18, 28-32). While many model systems for T6S contain multiple loci encoding different T6S machines, *Proteus mirabilis* strain BB2000 is one of many less-studied organisms with a single T6S system and single alleles for TssB (BB2000_0821) and TssC (BB2000_0820) (33, 34). We have previously reported that a point mutation in *BB2000_0821* results in the blocked export of proteins belonging to the Ids recognition system (35). This mutation lies in an unstructured region of the TssB monomer that was previously unexamined. Here we show that the identified point mutation disrupts global T6S export function. Based on published structures of the TssB-TssC sheath in other organisms, we mapped the point mutation to a region involved in sheath formation (25-27). We specifically demonstrate that this point mutation inhibits sheath polymerization and that this is the likely mechanism for disrupted T6S export. Since this single point mutation disrupts global T6S function without disrupting the structure of the operon, and likely the core membrane components, we posit that the mutation provides a valuable tool for studying T6S-associated phenotypes in multiple organisms.

## Materials and Methods

### Bacterial strains and media

Strains and plasmids used in this study are described in Table 1. *P. mirabilis* strains were maintained on LSW^-^ agar (36). CM55 blood agar base agar (Oxoid, Basingstoke, England) was used for swarm-permissive nutrient plates. Overnight cultures of all strains were grown in LB broth under aerobic conditions at 37°C. Antibiotics used were: 35 microgram/milliliter (μg/ml) kanamycin, 50 μg/ml chloramphenicol, 15 μg/ml tetracycline, and 25 μg/ml streptomycin. Kanamycin was added for plasmid maintenance where appropriate.

**Table 1.** Strains used in this study.

| Strain | Notes | Source |
| --- | --- | --- |
| HI4320 | Wild-type strain | (37, 38) |
| $\Delta ids::tssB_{wt}$ | This strain, derived from the *P. mirabilis* wild-type strain BB2000, produces TssB<sub>wt</sub> and none of the Ids proteins. | (39) |
| $icmF^*::tssB_{wt}$ | This strain was previously reported as strain *tssN\**. It contains a transposon insertion in gene *BB2000_0808*, which encodes the TssM/IcmF homolog in *P. mirabilis*. | (34) |
| $\Delta ids::tssB_{mut}$ | This strain was previously reported as strain CCS05. It produces the TssB<sub>L32R</sub> mutant variant and is deficient in T6S-mediated transport. | (35) |
| $\Delta ids::tssB_{wt}$ carrying (c.) pLW101 | This strain carries a derivative of the low-copy plasmid pIdsBB (39), which encodes the entire *ids* operon under control of its native promoter, and in which a FLAG epitope (N-DYKDDDDK-C) was inserted immediately before the *idsA* stop codon (34). This strain was used for the LC-MS/MS studies. | (34) |
| $\Delta ids::tssB_{mut}$ c. pLW101 | This strain carries pLW101. This strain was used for the LC-MS/MS studies. | (35) |
| $\Delta ids::tssB_{wt}-sfGFP$ | This strain produces TssB<sub>wt</sub> fused to sfGFP. | This study |
| $\Delta ids::tssB_{mut}-sfGFP$ | This strain produces TssB<sub>L32R</sub> fused to sfGFP. | This study |
| $icmF^*::tssB_{wt}-sfGFP$ | This $icmF^*::tssB_{wt}$-derived strain produces TssB<sub>wt</sub> fused to sfGFP. | This study |
| $\Delta ids::tssB_{wt}$ c. pIdsBB-ΔE | This strain was previously reported as strain CCS06. It produces BB2000-derived IdsD from a pIdsBB (39)-derived plasmid, but not IdsE. It also produces TssB<sub>wt</sub>. | (35) |
| $\Delta ids::tssB_{mut}$ c. pIdsBB-$\Delta E$ | This strain produces BB2000-derived IdsD from a pIdsBB (39)-derived plasmid, but not IdsE. It also produces TssB<sub>L32R</sub>. | (35) |
| $\Delta ids::tssB_{wt-sfgfp}$ c. pIdsBB-$\Delta E$ | This strain produces BB2000-derived IdsD from a pIdsBB (39)-derived plasmid, but not IdsE. It also produces TssB<sub>wt</sub> fused to sfGFP. | This study |
| $\Delta ids::tssB_{mut-sfgfp}$ c. pIdsBB-$\Delta E$ | This strain produces BB2000-derived IdsD from a pIdsBB (39)-derived plasmid, but not IdsE. It also produces TssB<sub>L32R</sub> fused to sfGFP. | This study |
| S17λpir | *E. coli* mating strain for moving plasmids into *P. mirabilis* | (40) |
| SM10λpir | *E. coli* mating strain for moving suicide vector pKNG101 (41) into *P. mirabilis* | (42) |

### Construction of strains Δ*ids::tssB_wt_-sfgfp* and Δ*ids::tssB_mut_-sfgfp*

BB2000-derived Δ*ids*:*:tssB_wt_-sfgfp* and Δ*ids*::*tssB_mut_-sfgfp* were constructed using 5’-GATTAGCAGCAATATCGAGCTC-3’ as the forward and 5’-GCGCGCTCTAGACCTTAAGTTAAACCAAATATAGCTG-3’ as the reverse primer to amplify *BB2000_0821,* encoding the *P. mirabilis* TssB homolog, and its upstream region from genomic DNA of the wild-type strain BB2000 (36) or of the BB2000-derived mutant strain Δ*ids*::*tssB_mut_* (35). The polymerase chain reaction (PCR) product was digested with DpnI (New England Biolabs, Ipswich, MA). Overlap extension PCR (43) was used to combine the PCR product and a gBlock (Integrated DNA Technologies, Inc., Coralville, IA) containing the gene encoding sfGFP (gBlock sequence: 5’CGCGGGCCCGGTATTACCCCATAAATAGTGCTCATGGTCTTGTTTAACATTTTCTG AATAGTTAAACATTTTTACAGGGTTTTCTGTTGGAGAGTATGGCAAACGTAATAAGA AACGCGGTGCTGTTAACCCTAAATAACGGGAATCTTCTGCTTCACGCAGTGAGCGCC ATTTAGTGTGTGCAGGGCCTTCAAAAACAGATTTTAGATCTTTAATTGCCGGCAGTT CAGCGTAGCTATTAATACCAAAGAAATTAGGTGAAACAGATGATAGGAATGGCGCG TGAGCCATTGCACCAACAGTACTAACATACTGCATTAACTTCATATCTGGCGCAGTA TTGTTAAAGGCGTAGTTACCAATAACTGTTGCAACAGGTTCACCACCAAATTGGCCG TATCCTGAAGAGTAGACGTGTTGATAGAAGCCTGATTGAACAATTTCTGGAGAAAAT TCAAAATCTTCTAATAACTCTTCTTTAGTTGCATGAAGAATATTGATCTTAATATTTT CTCTAAAATCAGTGCGATCAACGAGTAATTTTAAAGAACGCCATGAAGCTTCAATTT CTTGAAATTTAGGCGCGTGAAGAATTTCGTCAACTTGAGTACTTAGTTTATTATCAA GTTCAACAAGCATTTTATCGATTAATAATCGATTGATCTGTTGGTCTTCAGATTCACT GGCGAAAATATTACTAATAAAAGCAGCAACACCTTGTTTGGCAATATCGTATGCTTC TGTTTCTGGAGACATACGTGATTGAGCCATAATTTCATCAAGTAAAGAGCCTGTAGA TGCGGGGGCTTGTTGCTCTTGAGCTTCTGCATTTAGTGACATGAAATAATCCTCTATA AACATTATTTGTAGAGCTCATCCATGCCATGTGTAATCCCAGCAGCAGTTACAAACT CAAGAAGGACCATGTGGTCACGCTTTTCGTTGGGATCTTTCGAAAGGACAGATTGTG TCGACAGGTAATGGTTGTCTGGTAAAAGGACAGGGCCATCGCCAATTGGAGTATTTT GTTGATAATGGTCTGCTAGTTGAACGGAACCATCTTCAACGTTGTGGCGAATTTTGA AGTTAGCTTTGATTCCATTCTTTTGTTTGTCTGCCGTGATGTATACATTGTGTGAGTT AAAGTTGTACTCGAGTTTGTGTCCGAGAATGTTTCCATCTTCTTTAAAATCAATACCT TTTAACTCGATACGATTAACAAGGGTATCACCTTCAAACTTGACTTCAGCACGCGTC TTGTAGGTCCCGTCATCTTTGAAAGATATAGTGCGTTCCTGTACATAACCTTCGGGC ATGGCACTCTTGAAAAAGTCATGCCGTTTCATGTGATCCGGATAACGGGAAAAGCAT TGAACACCATAGGTCAGAGTAGTGACAAGTGTTGGCCATGGAACAGGTAGTTTTCC AGTAGTGCAAATAAATTTAAGGGTGAGTTTTCCGTTTGTAGCATCACCTTCACCCTCT CCACGGACAGAAAATTTGTGCCCATTAACATCACCATCTAATTCAACAAGAATTGGG ACAACTCCAGTGAAAAGTTCTTCTCCTTTGCTTCCTCCTCCAGCAGCAGCTTTTTGAT TAGCAGCAATATCGAGCTC-3’). 5’CGCGGGCCCGGTATTACCCCATAAATAGTGC-3’ and 5’-GCGCGCTCTAGACCTTAAGTTAAACCAAATATAGCTG -3’ were used as the forward and reverse primers, respectively. Restriction digestion with ApaI and XbaI (New England Biolabs, Ipswich, MA) was used to introduce the overlap extension PCR product into the suicide vector pKNG101 (41). The resulting vectors pCS48a (*tssB*_mut_-*sfgfp*) and pCS48b (*tssB_wt_-sfgfp*) were introduced into the mating strain *E. coli* SM10λpir (42) and then mated into BB2000-derived Δ*ids* (39). The resultant matings were subjected to antibiotic selection using tetracycline and streptomycin on LSW^-^ agar. Candidate strains were subjected to sucrose counter-selection to select for clones that integrated the target DNA sequence into the chromosome in exchange for the wild-type DNA sequence (44). 5’-GCCATCAACATCAAGTACTTTG-3’ and 5’-CATGAGCAGTCCAAATTGATC-3’ were used as the forward and reverse primers, respectively, to amplify the exchanged chromosomal regions by colony PCR. PCR reactions were purified and the entire exchanged region was sequenced to confirm strains. Sequencing was performed by GENEWIZ, Inc. (South Plainfield, NJ).

### Trichloroacetic acid precipitations, SDS-PAGE, and LC-MS/MS

Trichloroacetic acid precipitations were performed as previously described (34). Gel fragments corresponding to molecular weights of approximately 10 to 20, 20 to 40, 40 to 60 and 60 to 250 kilodaltons (kDa) were excised and subjected to liquid chromatography-mass spectrometry/mass spectrometry (LC-MS/MS), which was performed by the Taplin Biological Mass Spectrometry Facility (Harvard Medical School, Boston, MA).

### Colony expansion and co-swarm inhibition assays

Overnight cultures were normalized to an optical density at 600 nm (OD_600_) of 0.1 and swarm-permissive nutrient plates were inoculated with one microliter (μl) of normalized culture. Plates were incubated at 37 °C for 16 hours, and radii of actively migrating swarms were measured. Additionally, widths of individual swarm rings within the swarm colonies were recorded.

For co-swarm inhibition assays, strains were processed as described and mixed at a ratio of 1:1 where indicated. Swarm-permissive nutrient plates supplemented with Coomassie Blue (20 μg/ml) and Congo Red (40 μg/ml) were inoculated with one μl of normalized culture/culture mixes. Plates were incubated at 37 °C until boundary formation was visible by eye.

### Modeling of TssB_wt_ and TssB_L32R_

Swiss-Model (45-48) was used for all modeling, and the atomic model, PDB ID: 3j9g (from *<u>http://www.rcsb.org</u>* (25, 49, 50)) was used as a template. Resulting .pdb files were modified in PyMOL v1.8.4.1 (51).

### Microscopy

One millimeter thick swarm-permissive agar pads were inoculated directly from overnight cultures. The agar pads were incubated at 37 °C in a modified humidity chamber. After 4.5 – 5.5 hours, the pads were imaged by phase contrast as well as epifluorescence microscopy using a Leica DM5500B (Leica Microsystems, Buffalo Grove, IL) and a CoolSnap HQ^2^ cooled CCD camera (Photometrics, Tucson, AZ). MetaMorph version 7.8.0.0 (Molecular Devices, Sunnyvale, CA) was used for image acquisition.

### Western blotting

To test for the production of the sfGFP-fused proteins, cells were isolated from swarm-permissive nutrient plates supplemented with chloramphenicol after 16-20 hours, resuspended in 5 ml LB broth, and normalized to an OD_600_ of 1. Cells were pelleted by centrifugation and the pellet was resuspended in sample buffer and boiled. Samples were separated by gel electrophoresis using 12% Tris-tricine polyacrylamide gels, transferred onto 0.45 micrometer nitrocellulose membranes, and probed with polyclonal rabbit anti-GFP (1:4000, ThermoFisher Scientific, Waltham, MA) or mouse anti-σ^70^ (1:1000, BioLegend, San Diego,CA, catalog number 663202) followed by goat anti-rabbit or goat anti-mouse conjugated to horseradish peroxidase. The polyclonal rabbit anti-GFP antibody was supplied at a concentration of 2 mg/ml (Thermo Fisher Scientific, Waltham, MA, catalog number A-11122, immunogen: GFP from jellyfish *Aequorea Victoria*). The goat anti-mouse secondary antibody was used at a 1:5000 dilution (KPL, Inc., Gaithersburg, MD, catalog number 214-1806) as was the goat anti-rabbit secondary (KPL, Inc., Gaithersburg, MD, catalog number 074-1506). A Precision Plus Protein^TM^ Dual Color Standards molecular size marker (Bio-Rad Laboratories, Inc., Hercules, CA) was included as a size control. Membranes were developed with the Immun-Star HRP Substrate Kit (Bio-Rad Laboratories, Hercules, CA) and visualized using a Chemidoc (Bio-Rad Laboratories, Hercules, CA). TIFF images were exported and figures were made in Adobe Illustrator (Adobe Systems, San Jose, CA).

## Results

### The tssB mutant strain does not export self-recognition proteins

Several self-recognition proteins in *P. mirabilis* strain BB2000 are exported by its T6S system; these include the Hcp (Pfam family PF05638) homologs IdsA and IdrA, the VgrG (Pfam family PF05954) homologs IdsB and IdrB, and the self-identity protein IdsD (34). We previously reported that a strain expressing a point mutation (L32R) from gene *BB2000_0821*, which encodes the *P. mirabilis* TssB homolog, results in the lack of IdsA, IdsB, and IdsD export (35). To determine whether global T6S export was disrupted or whether the defect was specific to proteins belonging to the Ids system, we examined the Idr-specific secretion profile of the mutant strain. We collected supernatants of liquid-grown cells producing either TssB_wt_ or TssB_L32R_. These supernatants were subjected to trichloroacetic acid precipitations followed by LC-MS/MS analysis. We identified peptides belonging to IdrA (Hcp), IdrB (VgrG) and the putative Idr effector IdrD in the supernatants of the wild type, but not of the mutant strain (Table 2).

**Table 2.** Idr-specific LC-MS/MS results of supernatant fractions from strains producing TssB_wt_ or TssB_L32R_

| Strain | TssB variant produced | Protein | Predicted size (kDa) | Gel slice (approx. kDa range) | No. of unique peptides | No. of total peptides | Coverage (%) |
| --- | --- | --- | --- | --- | --- | --- | --- |
| *Δids::tssB<sub>wt</sub>*<br>c. pLW101 | TssB<sub>wt</sub> | σ<sup>70</sup> | 71.11 | 60 - 250* | 0 | 0 | 0 |
|  |  |  |  | 40 - 60 | 0 | 0 | 0 |
|  |  |  |  | 20 - 40 | 2 | 2 | 4.37 |
|  |  |  |  | 10 - 20 | 0 | 0 | 0 |
|  |  | IdrA | 18.99 | 60 - 250<sup>1</sup> | 2 | 3 | 13.95 |
|  |  |  |  | 40 - 60 | 0 | 0 | 0 |
|  |  |  |  | 20 - 40<sup>2</sup> | 8 | 29 | 55.81 |
|  |  |  |  | 10 - 20*<sup>3</sup> | 4 | 21 | 26.74 |
|  |  | IdrB | 80.23 | 60 - 250* | 4 | 4 | 7.26 |
|  |  |  |  | 40 - 60 | 0 | 0 | 0 |
|  |  |  |  | 20 - 40 | 2 | 2 | 4.52 |
|  |  |  |  | 10 - 20 | 2 | 2 | 6.58 |
|  |  | IdrD | 178.29 | 60 - 250* | 2 | 2 | 1.33 |
|  |  |  |  | 40 - 60 | 2 | 2 | 1.27 |
|  |  |  |  | 20 - 40 | 7 | 10 | 3.16 |
|  |  |  |  | 10 - 20 | 8 | 59 | 3.73 |
| *Δids::tssB<sub>mut</sub>*<br>c. pLW101 | TssB<sub>L32R</sub> | $\sigma^{70}$ | 71.11 | 60 - 250* | 0 | 0 | 0 |
|  |  |  |  | 40 - 60 | 0 | 0 | 0 |
|  |  |  |  | 20 - 40 | 0 | 0 | 0 |
|  |  |  |  | 10 - 20 | 0 | 0 | 0 |
|  |  | IdrA | 18.99 | 60 - 250 | 0 | 0 | 0 |
|  |  |  |  | 40 - 60 | 0 | 0 | 0 |
|  |  |  |  | 20 - 40 | 0 | 0 | 0 |
|  |  |  |  | 10 - 20*<sup>1</sup> | 2 | 2 | 13.95 |
|  |  | IdrB | 80.23 | 60 - 250* | 0 | 0 | 0 |
|  |  |  |  | 40 - 60 | 0 | 0 | 0 |
|  |  |  |  | 20 - 40 | 0 | 0 | 0 |
|  |  |  |  | 10 - 20 | 0 | 0 | 0 |
|  |  | IdrD | 178.29 | 60 - 250* | 0 | 0 | 0 |
|  |  |  |  | 40 - 60 | 0 | 0 | 0 |
|  |  |  |  | 20 - 40 | 0 | 0 | 0 |
|  |  |  |  | 10 - 20 | 0 | 0 | 0 |

\* Expected kDa range according to molecular weight
<sup>1</sup> Given the high degree of sequence similarity between the Hcp homologs IdsA and IdrA, all

To confirm these secretion profiles, we employed an *in vivo* assay for Idr self-recognition activity. The Idr proteins contribute to inter-strain competitions during swarm migration (34), which is a highly coordinated, flagella-based social motility in *P. mirabilis* (52-54). Briefly, the assay consists of mixing two different strains of *P. mirabilis* at a 1:1 ratio and allowing for swarm migration; at the completion of swarm migration, one strain dominates over the other and thus occupies the outer edges of the swarm colony (34). The dominating strain can be identified by swarm boundary formation assays in which two genetically identical strains merge to form a single larger swarm, while two genetically distinct strains remain separate and form a visible boundary (Fig 1A) (39, 55-57). *P. mirabilis* strain BB2000 dominates over the genetically distinct strain HI4320, permitting BB2000 to occupy the outer edges of a mixed swarm colony, which thus merges with the swarm colony of BB2000 and forms a boundary with the swarm colony of HI4320 (34). Absence of Idr proteins or of T6S in BB2000 allows for HI4320 to dominate the mixed swarm instead (34). Loss of the Ids proteins in BB2000 has no effect (34). We therefore performed these assays using BB2000-derived strains that lack the Ids proteins (Δ*ids*) and produce either TssB_wt_ or the mutant TssB_L32R_. As a control, we used the BB2000-derived strain, *icmF\**, which contains a disruption in the gene encoding the T6S core membrane component TssM/IcmF (Pfam family PF12790); this mutant strain is defective for T6S function (33, 34). As expected, Δ*ids* producing TssB_wt_ dominated over strain HI4320, and conversely, strain HI4320 dominated over *icmF\** (Fig 1B). Δ*ids* producing the mutant TssB_L32R_ did not dominate over HI4320 (Fig 1B), indicating that Idr-dependent inter-strain competition was disrupted in this strain. As co-swarm inhibition of HI4320 by BB2000-derived strains is dependent on the export of Idr self-recognition proteins, we conclude that a strain producing TssB_L32R_ is defective in the export of Idr proteins. Together, the secretion profile and swarm assays support that this mutation (TssB_L32R_) causes a global loss in T6S export activity.

**Figure 1.**
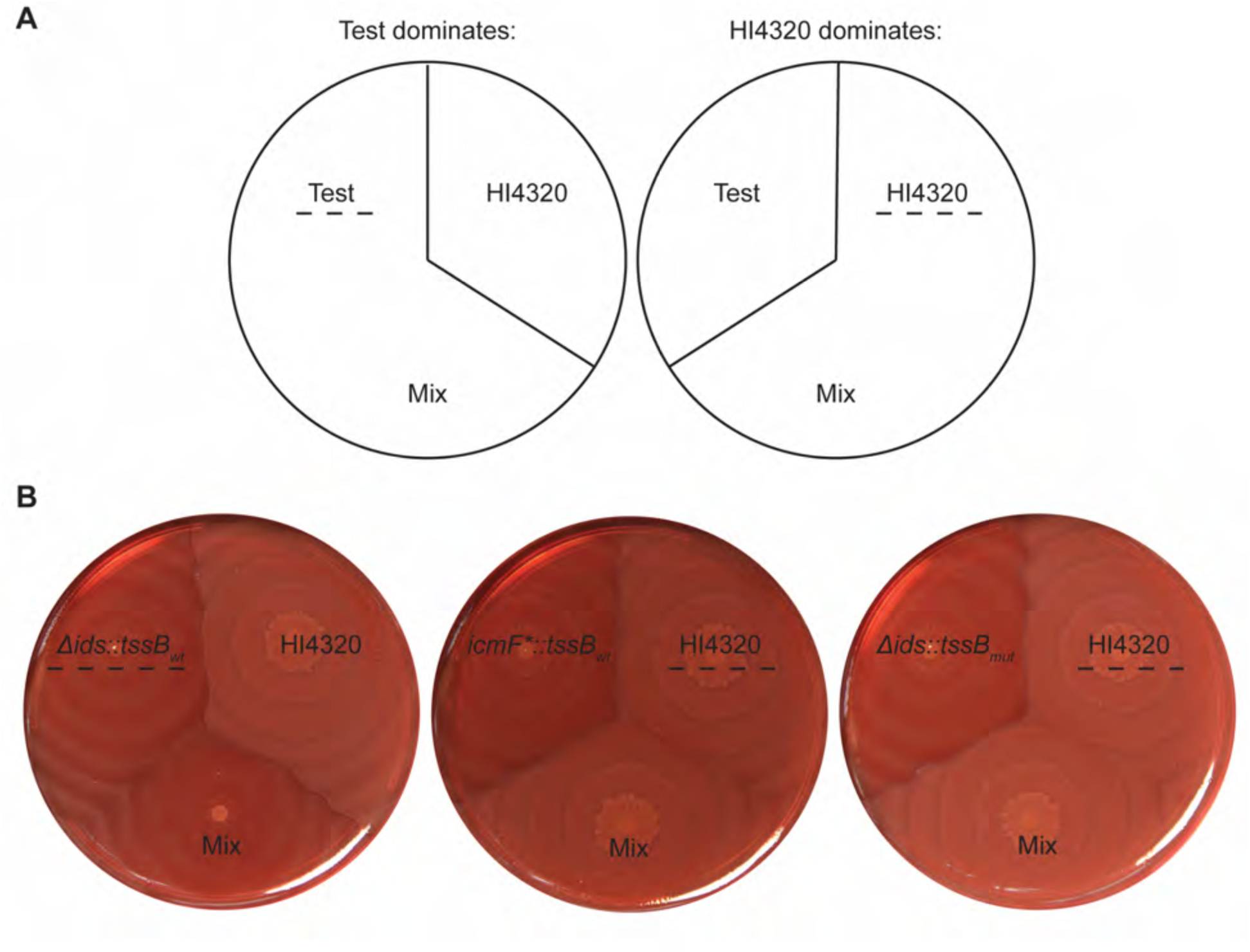
The L32R mutation disrupts Idr function. (A) Schematic of the Idr-dependent co-swarm assay. Strains are inoculated either as monocultures or as 1:1 mixed cultures onto swarm-permissive media as previously described (34). The dominating strain in the mixed culture will occupy the outer swarm edges; boundary formation assays (39, 55-57) are then used to determine the identity of the dominating strain. The test strain dominated the mixed swarm if the mixed swarm colony forms a visible boundary with strain HI4320. Conversely, strain HI4320 dominated the mixed swarm if the mixed swarm does not form a boundary with strain HI4320. The dominating strain is indicated by a dashed line. (B) The Idr-dependent co-swarm assay using the indicated strains. BB2000-derived Δ*ids::tssB_wt_* produces TssB_wt_; it lacks the entire *ids* operon (39). BB2000-derived *icmF*::tssB_wt_* produces TssB_wt_; it contains a chromosomal transposon insertion in the gene encoding the core T6S membrane component, TssM/IcmF (34). Δ*ids*::*tssB_mut_* (35) produces the mutant variant TssB_L32R_.

### The L32R mutation in the TssB homolog prohibits sheath formation

To understand the disrupted T6S export, we modeled the *P. mirabilis* BB2000 T6S sheath structure on a published structure of the *Vibrio cholerae* T6S sheath (25) (Fig 2). We then mapped the L32R point mutation onto this *P. mirabilis* model structure (Fig 2). The point mutation causes a hydrophobic leucine at position 32 to be substituted with a positively charged arginine, which has a larger side chain and different electrochemical properties (Fig 2A). Amino acid 32 maps to an unstructured region between the first two beta sheets of TssB (Figs 2B, 2C) (25, 26). The first beta sheet is involved in interactions between TssB-TssC protomers that hold together the strands of the sheath helix, while the second beta sheet is involved in TssB-TssC interactions within individual protomers (Figs 2B, 2C) (25, 26). Given the position within the T6S sheath, we hypothesized that the L32R mutation interferes with sheath formation.

**Figure 2.**
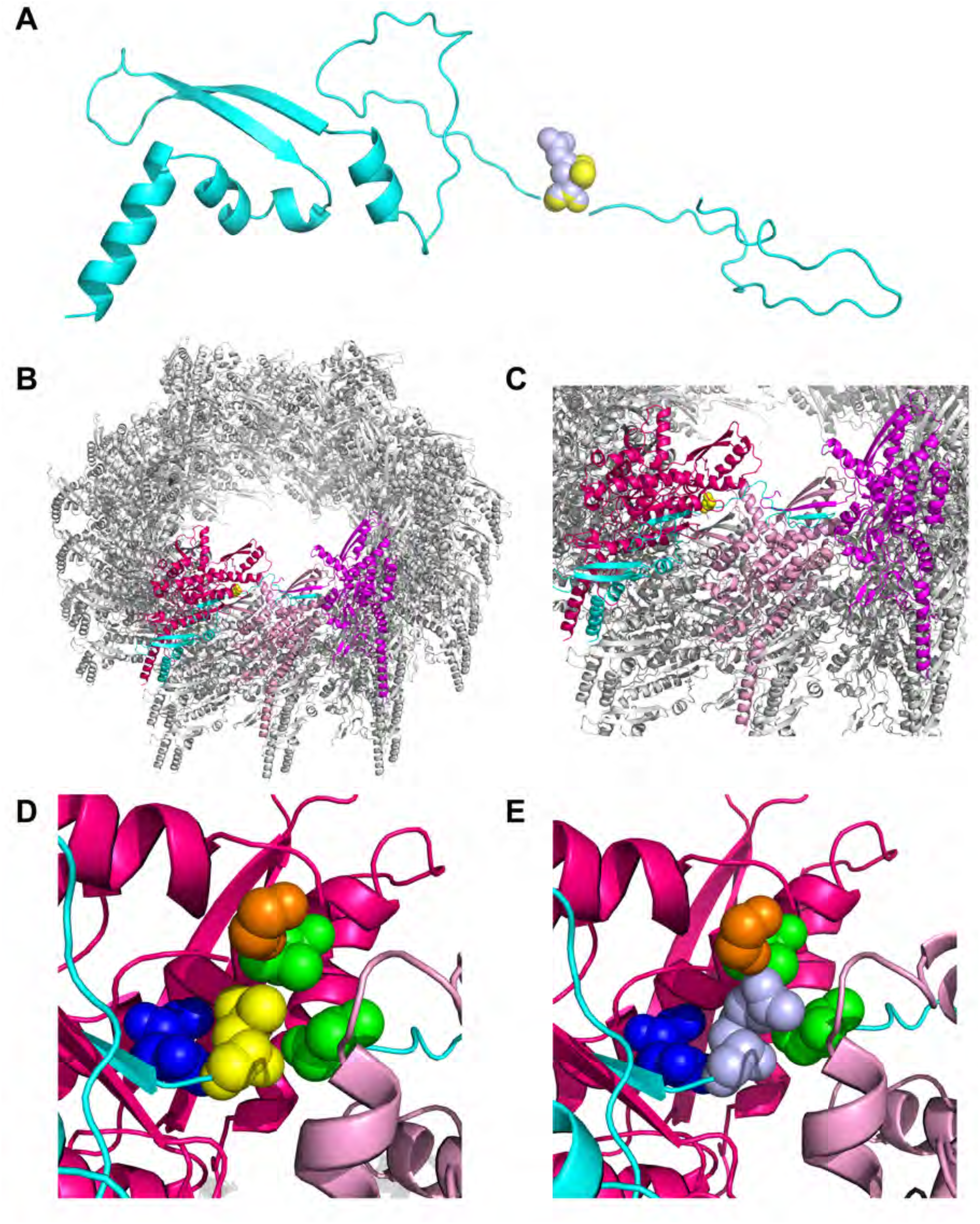
The L32R mutation on a model of the *P. mirabilis* BB2000 T6S sheath. (A) Cartoon representation of the *P. mirabilis* BB2000 TssB monomer (cyan) modeled after the *V. cholerae* TssB homolog (25). The wild-type TssB variant has a leucine (yellow) at position 32. The mutant variant TssB_L32R_ has an arginine (light blue) at position 32. (B, C) Cartoon representation of a portion of the wild-type *P. mirabilis* BB2000 T6S sheath containing TssB_wt_ and the *P. mirabilis* BB2000 TssC homolog (BB2000_0820) at two magnifications. It was modeled after the *V. cholerae* contracted sheath (25). In this model, one complete TssB-TssC protomer is highlighted, with TssB shown in cyan and TssC shown in pink. Two additional TssC monomers are highlighted in magenta and light pink. The wild-type TssB variant has a leucine (yellow) at position 32, which we have mapped to a predicted unstructured region between two beta sheets. Both beta sheets are thought to be involved in making contacts to TssC monomers (25, 26). (D, E) Magnified view of residue 32 and the neighboring residues (glycine in orange, valine in green, and arginine in dark blue). (D) Wild-type leucine in yellow. (E) Mutant arginine in light blue. Swiss-Model (45-48) was used for all modeling, and the atomic model 3j9g (from *http://www.rcsb.org* (25, 49, 50)) was used as a template. Resulting .pdb files were modified in PyMOL v1.8.4.1 (51).

We introduced the mutation into a Δ*ids* strain producing TssB fused to sfGFP to directly observe sheath formation *in vivo*. To validate the functionality of this tool in *P. mirabilis* strain BB2000, we examined the ability of the TssB-sfGFP variants to export self-recognition proteins using established assays. We subjected Δ*ids*::*tssB_wt_-sfgfp* and Δ*ids*::*tssB_mut_-sfgfp* to Idr-dependent inter-strain competitions with strain HI4320 as described above. Surprisingly, HI4320 dominated over both strains (Fig 3A), which suggested that T6S export is impaired in both strains. A more quantitative assay for T6S function is a swarm expansion assay using the Ids proteins; in this assay, a reduced swarm colony radius indicates that the self-identity determinant IdsD has been exported and that T6S is functional (35). As previously reported (35), the presence of wild-type TssB results in a reduced swarm colony radius, while disrupted T6S function due to the presence of TssB_L32R_ leads to a larger swarm colony radius (Fig 3B). We examined the strain producing TssB_wt_-sfGFP and found that its swarm colony radius was modestly increased in comparison to that of the strain producing untagged TssB_wt_ (Fig 3B); therefore, fusion of sfGFP to TssB_wt_ caused a modest reduction in T6S function. Interestingly, the reduced functionality of TssB_wt_-sfGFP in comparison to TssB_wt_ was sufficient to prevent a BB2000-derived strain to outcompete HI4320 in the *idr*-specific co-swarm assay (Fig 3A), further supporting earlier observations of distinct functions for Ids and Idr proteins (34). As expected, the strain producing TssB_L32R_-sfGFP exhibited an even larger swarm colony radius, similar to that of a strain producing untagged TssB_L32R_ (Fig 3B). This observation demonstrates a lack of T6S export (35) and thus confirms that strains producing TssB_L32R_ do no export the T6S substrate, IdsD.

**Figure 3.**
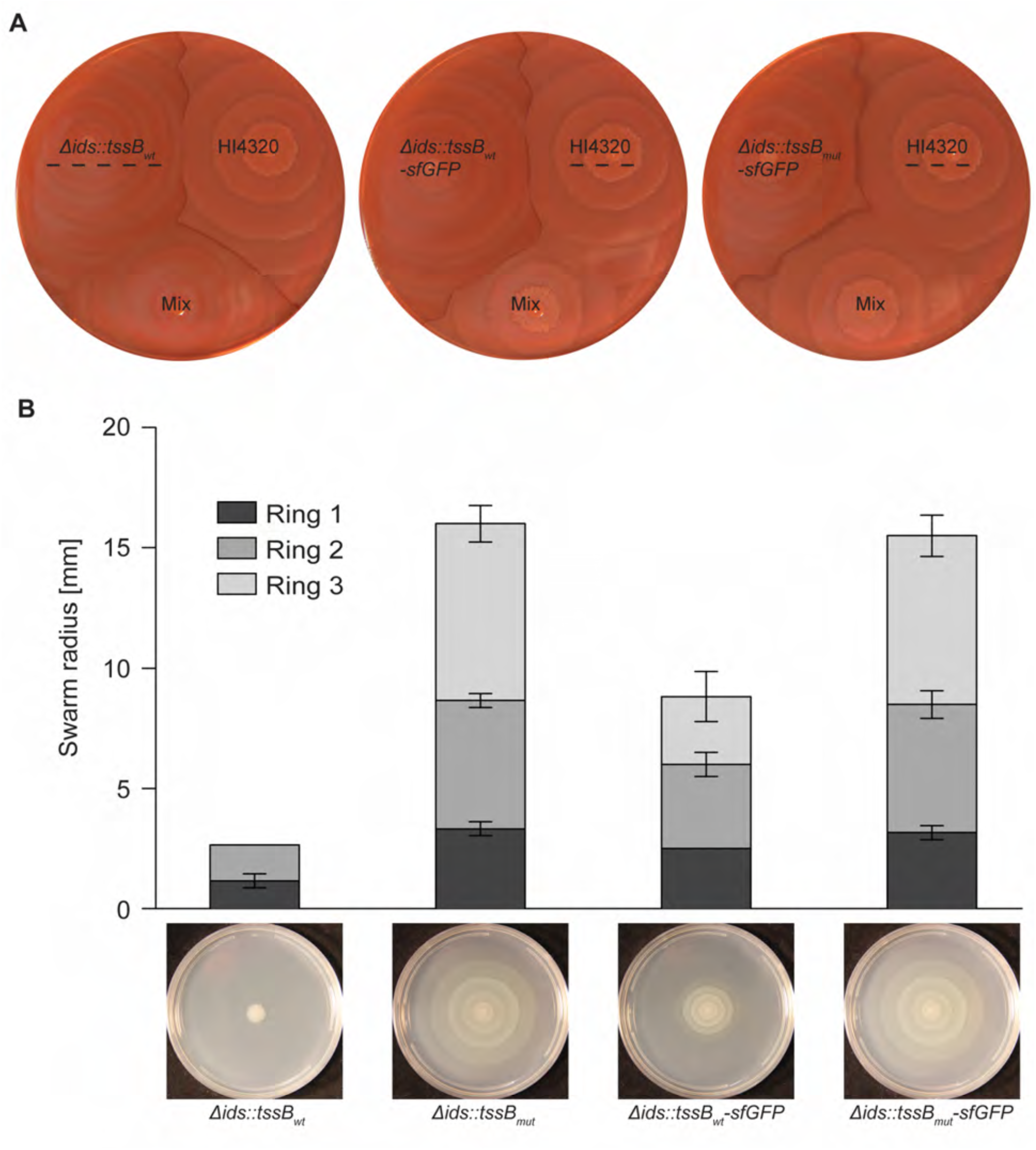
The TssB_wt_-sfGFP fusion is partially functional in *P. mirabilis*. (A) Idr-dependent co-swarm assays were performed as described in Fig 1. Δ*ids::tssB_wt_* produces TssB_wt_; Δ*ids*::*tssB_wt_-sfgfp* produces wild-type TssB fused to sfGFP (TssB_wt_-sfGFP); and Δ*ids*::*tssB_mut-_sfgfp* produces the L32R mutant variant of TssB fused to sfGFP (TssB_L32R-_sfGFP). The dominating strain is indicated with a dashed line on all plates. (B) Swarm assay in which a reduced swarm colony radius denotes that IdsD has been exported and transferred to an adjacent cell, where it remained unbound due to the lack of its binding partner IdsE (35, 58). Wild-type T6S function results in a reduced swarm colony radius; disrupted T6S function results in larger swarm colony radii. Shown is the colony expansion after 16 hours on swarm-permissive agar surfaces of monoclonal Δ*ids*-derived swarms producing IdsD and lacking IdsE (35). Strains contain the indicated *tssB* alleles. Widths of individual swarm rings within a swarm colony are marked by different shades. Representative images of swarm colonies after 24 hours are shown below the graph. N = 3, error bars show standard deviations of individual swarm ring widths.

We next employed the TssB-sfGFP tool to determine the ability of the mutant TssB variant to support T6S sheath formation, thereby testing the hypothesis that TssB_L32R_ does not support sheath formation. Fluorescence associated with TssB_wt_-sfGFP appears in multiple discrete (short or elongated) structures throughout cells on swarm-permissive agar (Fig 4A). This fluorescence pattern is consistent with previous reports for TssB-sfGFP (6, 59). In cells producing the mutant fusion protein, diffuse fluorescence signal was present, and no elongated structures or puncta were observed (Fig 4A). Therefore, the discrete structures, thought to be sheaths, did not form. To further examine whether the discrete fluorescence was likely representative of assembled T6S sheaths, we produced TssB_wt_-sfGFP in the *icmF\** strain, which lacks the membrane-associated complex of the T6S machinery and which we thus predicted to lack sheaths. Again we observed that fluorescence was diffuse in these cells (Fig 4A), indicating the likely absence of sheath formation.

**Figure 4.**
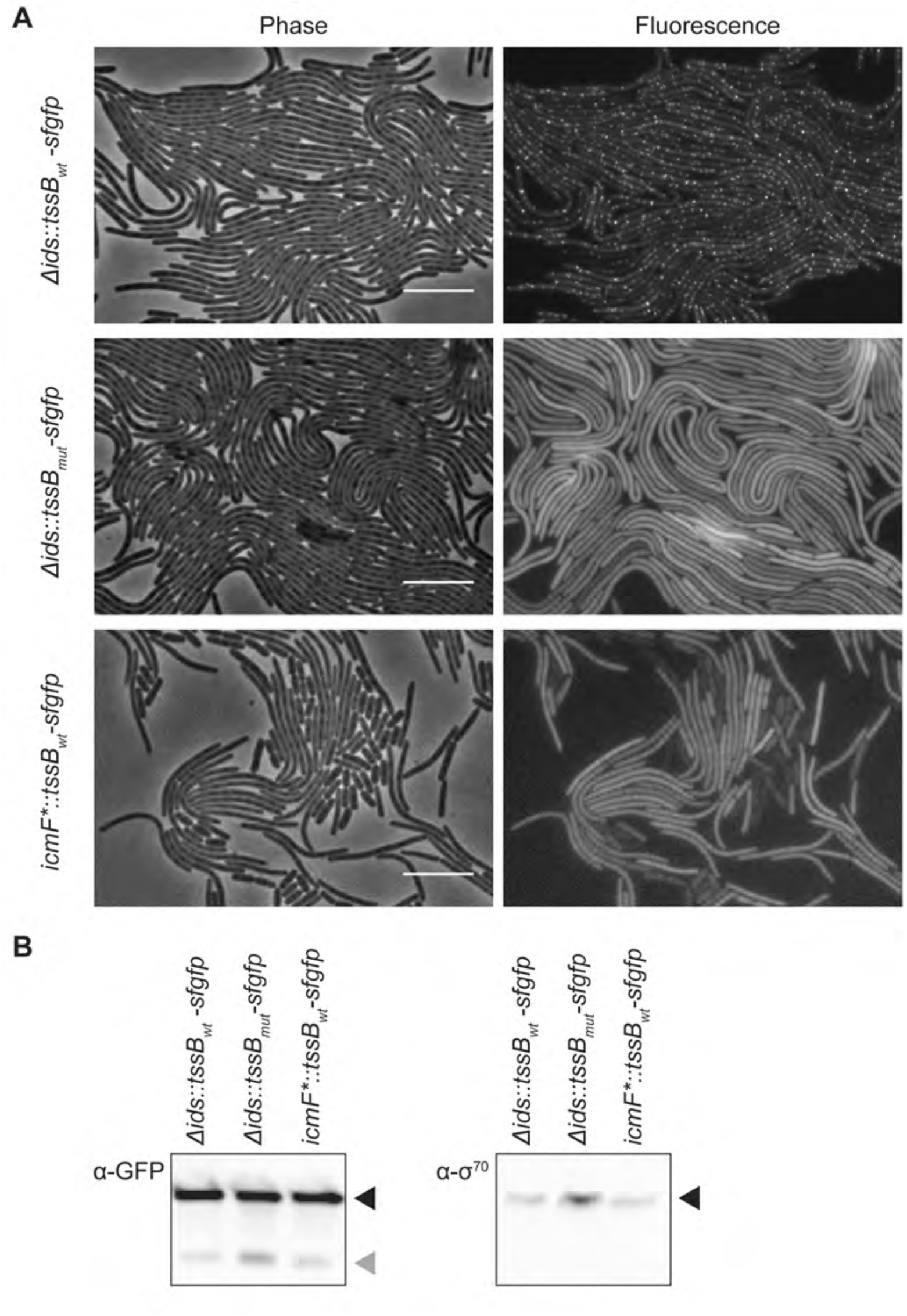
Sheath formation is inhibited by the L32R mutation. (A) Swarm agar pads were inoculated with indicated strains and incubated at 37 °C in a humidity chamber. After 4.5 – 5.5 hours, agar pads were imaged using phase contrast and epifluorescence microscopy to visualize cell bodies and TssB-sfGFP variants, respectively. BB2000-derived Δ*ids*::*tssB_wt_-sfgfp* produces wild-type TssB fused to sfGFP (TssB_wt_-sfGFP); it lacks the entire *ids* operon (39). Δ*ids*::*tssB_mut_-sfgfp* produces the L32R mutant variant of TssB fused to sfGFP (TssB_L32R_-sfGFP). *icmF*::tssB_wt_-sfgfp* produces TssB_wt_-sfGFP; it is derived from BB2000 and contains a chromosomal transposon insertion in the gene encoding the core T6S membrane component, TssM/IcmF (34). Scale bars, 10 μm. (B) Whole cell extracts from swarming colonies of Δ*ids*::*tssB_wt_-sfgfp*, Δ*ids*::*tssB_mut_-sfgfp* and *icmF*::tssB_wt_-sfgfp* were collected after 16-20 hours on swarm-permissive plates. Samples were analyzed using western blot analysis and probed with an anti-GFP antibody to detect TssB-sfGFP and anti-σ^70^ as a loading control. Bands corresponding to the sizes of TssB-sfGFP and σ^70^ are indicated with black arrowheads, while bands corresponding to the size of monomeric sfGFP are indicated with a grey arrowhead. A negative control sample of a swarm colony not producing any TssB-sfGFP fusion can be found in Fig S1.

A possible explanation for the diffuse fluorescence pattern in a strain producing TssB_L32R_-sfGFP could be that the mutant TssB is less stable than the wild-type variant, which could result in the cleavage of sfGFP from the fusion protein and ultimately a diffuse signal. We examined whole cell extracts that were collected from swarming colonies and then subjected these extracts to western blotting followed by incubation with anti-GFP antibodies. We found no striking differences among the strains producing wild-type or mutant TssB-sfGFP (Fig 4B). For both strains, we observed a dominant band corresponding to the size of the TssB-sfGFP fusion protein and a fainter band corresponding to the size of monomeric sfGFP (Fig 4B). This result suggests that while cleavage of the TssB-sfGFP fusion occurs, it does so for both the wild-type and mutant TssB-sfGFP variants. We conclude that cleavage of the sfGFP does not account for the differences in fluorescence patterns of these strains. Therefore, the L32R mutation in TssB is sufficient to prohibit sheath formation in *P. mirabilis*.

## Discussion

Here we have described the critical contribution of a single residue in TssB for sheath formation and global T6S export activity. We have shown that a L32R mutation in the *P. mirabilis* TssB variant inhibits sheath assembly (Fig 4A) without altering relative amounts of this protein (Fig 4B). The mutated leucine residue in the *P. mirabilis* variant corresponds to amino acid 26 in the Pfam model, T6SS_TssB (Pfam family PF05591) (24, 60-62). Leucine 26 is highly conserved in available sequences for Enterobacterales and is also well conserved across the majority of Gammaproteobacteria; the conserved three amino acid sequence amongst Enterobacterales is E/Q-L-P (63-65). Remarkably, the leucine residues maps to an unstructured region between the first two predicted beta sheets of the TssB protein (Fig 2). While the two beta sheets flanking this residue are predicted to be involved in interactions with separate TssC monomers in structural models of the sheath (Fig 2), the contribution of the unstructured region was less apparent (25, 26). Upon closer evaluation, we posit that this leucine residue might contribute to a hydrophobic pocket (Fig 2D). A conversion to arginine then likely leads to both steric and electrostatic conflicts within this pocket (Fig 2E). The hydrophobic pocket and the examined leucine residue within it might be critical for interactions between TssB and TssC in individual protomers or alternatively, for interprotomer interactions of TssB and TssC that hold together the strands of the sheath helix. More examination is needed to elucidate these physical interactions. Nonetheless, the observation that the L32R mutation results in diffuse TssB-sfGFP signal within the cytoplasm (Fig 4A) supports the crucial role of this leucine for sheath assembly.

Visualization of sheath formation and its subcellular localization has proven valuable in multiple bacteria. TssB-sfGFP fusion proteins have been previously reported for *Vibrio cholerae* (6) and *P. mirabilis* strain HI4320 (59); the homologous TssB-sfGFP was designed equivalently in *P. mirabilis* strain BB2000 and introduced to the native locus on the chromosome replacing the wild-type allele. This sfGFP fusion protein showed reduced T6S function as compared to the wild-type variant (Fig 3). As similar defects were not reported previously (6, 59), we posit that possible differences in the protein sequences and tertiary structures between species and strains might be accentuated by the addition of sfGFP. Alternatively, these discrepancies in T6S on the species and strain level might reflect differences in the sensitivity of the respective assays chosen to detect T6S function (Fig 3). Regardless, the reduced T6S function in *P. mirabilis* strain BB2000 producing TssB-sfGFP appears to allow for greater visualization of elongated sheaths before disassembly (Fig 4). Elongated sheaths were stable over the course of the observation. Similar delayed dynamics might also be true for other microorganisms with nearly identical TssB sequences and a single T6S machinery such as *Providencia sp*. By contrast, TssB-sfGFP-containing sheaths rapidly transition between elongated structures and single foci during imaging in *V. cholerae* (4-6). Stable, elongated TssB-sfGFP structures, such as those observed in *P. mirabilis* BB2000, could be utilized for subcellular co-localization assays using epifluorescence microscopy or for structural examination of the T6S machinery via cryo-electron microscopy.

In conclusion, the L to R mutation has the potential to be an impactful tool, especially in bacteria with a single T6S machinery. We have found that the L32R point mutation in the TssB homolog from *P. mirabilis* strain BB2000 allows for the disruption of the T6S sheath, which in turn causes global T6S malfunction. These results provide insight into the importance of the TssB N-terminal region for sheath assembly, which has not been fully investigated, and highlights a critical residue that was previously overlooked. These data also complement a previous detailed analysis of the TssB C-terminal region (28). Last, the L to R mutation abrogates T6S function without disrupting the overall *tss* operon structure and without removing entire proteins from cells. As such, potential secondary effects that could convolute experimental outcomes are minimized and questions regarding the subcellular localization of T6S-associated components and protein-protein interactions can be answered in a nearly native state. Here, we show that this previously underappreciated residue is indeed crucial for sheath assembly and raise new questions of how unstructured regions within TssB contribute to TssB-TssC interactions.

## Acknowledgments

We thank Dr. Thomas Bernhardt for the gift of the *sfgfp* sequence as well as Dr. Kenneth Skinner for helpful advice on the structural modeling. We also thank the members of Dr. Rachelle Gaudet’s research group, as well as the Gibbs research group, for thoughtful discussions.

